# Curcumin Analogues Trigger HMOX1-Mediated Ferroptosis to Halt Endometrial Cancer Growth

**DOI:** 10.1101/2025.11.18.689080

**Authors:** Xiangxiang Wu, Jamie L Padilla, Lane E Smith, Lavanya Goodla, Kimberly K. Leslie, Xiang Xue

## Abstract

**Background:** Advanced endometrial cancer remains challenging to treat due to limited therapeutic options and drug resistance. Ferroptosis, an iron-dependent form of cell death, offers a potential strategy for overcoming resistance. Curcumin analogues with improved bioavailability, such as HO-3867 and AKT-100, exhibit potent anti-cancer activity, but their mechanisms remain under-explored.

**Methods:** KLE, Hec50co and Ishikawa endometrial cancer cells were treated with HO-3867 or AKT-100. RNA sequencing, qPCR, and immunoblotting assessed ferroptosis-related gene expression, focusing on HMOX1. Intracellular iron and ROS were measured via FerroOrange and DCFH-DA staining. Cytotoxicity and colony formation were evaluated using CyQUANT assays, with pharmacological inhibitors (ZnPP, Liproxstatin-1, Z-VAD-FMK) and HMOX1 siRNA to dissect the roles of ferroptosis and apoptosis.

**Results:** Both analogues upregulated multiple ferroptosis genes, prominently HMOX1. AKT-100 increased intracellular iron and ROS levels. Inhibition of HMOX1, ferroptosis, or apoptosis partially rescued cell viability, while HMOX1 knockdown enhanced clonogenic growth, confirming its key role in AKT-100–mediated cytotoxicity.

**Conclusions:** HO-3867 and AKT-100 induce HMOX1–mediated ferroptosis and apoptosis, effectively suppressing endometrial cancer cell growth. These findings support the therapeutic potential of curcumin analogues and provide a foundation for *in vivo* studies targeting HMOX1 in endometrial cancer.

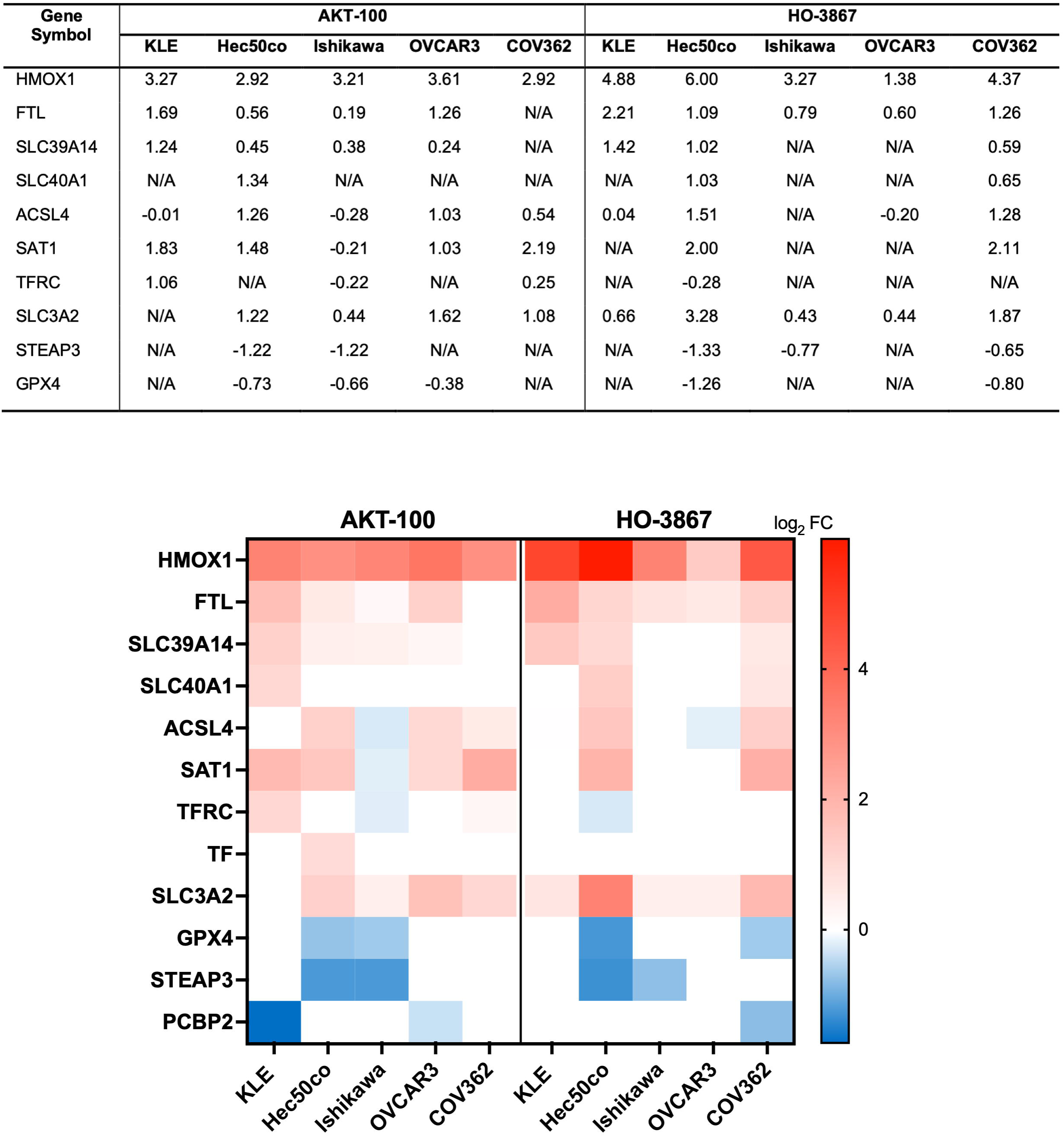

## Introduction

Endometrial cancer is the most common gynecologic malignancy in developed countries, and while early-stage disease is generally curable, advanced-stage or recurrent tumors are associated with poor outcomes, with 5-year survival rates as low as 0–18% (Siegel et al., 2022). Standard therapies—including surgery, radiotherapy, and chemotherapy—are often ineffective in metastatic or treatment-resistant cases (Carlson et al., 2014; Brasseur et al., 2022). Resistance to apoptosis (Mohammad RM et al., 2015), the primary pathway targeted by conventional treatments, remains a major obstacle, highlighting the urgent need to identify alternative mechanisms of tumor cell elimination and to develop novel therapeutics capable of overcoming drug resistance.

Ferroptosis, an iron-dependent, non-apoptotic form of regulated cell death, has emerged as a promising anticancer strategy (Dixon et al., 2012). Ferroptosis is characterized by the accumulation of intracellular labile iron, elevated reactive oxygen species (ROS), and lipid peroxidation, ultimately resulting in oxidative membrane damage and cell death (Mao C, et al. 2025). Unlike apoptosis, ferroptosis is less susceptible to resistance mechanisms in cancer cells, making it a particularly attractive target in therapy-refractory tumors (Zhang et al., 2022). Pharmacologic induction of ferroptosis has been shown to restore sensitivity to conventional therapies, reverse acquired drug resistance and enhance the efficacy of chemotherapeutics and targeted agents across multiple cancer types (Li et al., 2023; Ghoochani et al., 2021). Furthermore, ferroptosis-related genes have been proposed as prognostic biomarkers in endometrial cancer, offering both mechanistic insight and potential clinical utility (Liu et al., 2022).

Curcumin, a naturally occurring polyphenol derived from turmeric, has demonstrated anti-cancer effects, including induction of ferroptosis, but its poor bioavailability and rapid metabolism limit clinical application (Xin and Zhang, 2024). To overcome these limitations, bioavailable curcumin analogues such as HO-3867 (Smith et al., 2025) and AKT-100 (Williams et al., 2025) have been developed. These compounds exhibit potent cytotoxicity at low micromolar to nanomolar concentrations in endometrial tumor cells while sparing non-cancerous cells (Selvendiran et al., 2010). HO-3867 has been explored in limited preclinical studies (Rath KS et al., 2014), whereas AKT-100 remains largely uncharacterized, highlighting the need for mechanistic studies in endometrial cancer.

Heme oxygenase-1 (HMOX1) is a stress-inducible enzyme that catalyzes heme degradation, releasing free iron, carbon monoxide, and biliverdin (Soares MP et al., 2009). HMOX1 plays a dual role in cancer biology: while it can provide cytoprotective effects under oxidative stress, its overactivation can elevate the labile iron pool and reactive oxygen species (ROS), thereby sensitizing cells to ferroptosis (Sun et al., 2021). Dysregulation of HMOX1 has been associated with tumor progression, therapy resistance, and modulation of redox homeostasis, making it a compelling target for ferroptosis-based therapeutics (Zuo L et al., 2025). Induction of HMOX1 by curcumin analogues may thus integrate iron metabolism and ROS accumulation to trigger ferroptotic cell death, potentially complementing apoptotic pathways and enhancing overall tumor suppression.

In this study, we investigated how HO-3867 and AKT-100 modulate ferroptosis-related gene expression, with a particular focus on HMOX1, in endometrial cancer cells. Using a combination of pharmacologic inhibitors, genetic knockdown approaches, and functional assays, we evaluated the interplay between ferroptosis and apoptosis, intracellular iron dynamics, ROS generation, and clonogenic potential. This work aims to elucidate the mechanistic underpinnings of curcumin analogue–mediated tumor suppression and provide a robust preclinical rationale for targeting HMOX1–regulated ferroptosis as a therapeutic strategy in advanced and drug-resistant endometrial cancer.

## Materials and Methods

### Cell Culture

KLE cells (ATCC #CRL-1622) were cultured in RPMI-1640 medium supplemented with 20% fetal bovine serum (FBS), 1% penicillin–streptomycin, and 1% non-essential amino acids. Hec50co cells and Ishikawa cells were acquired form Dr. Erlio Gurpide, New York University, and both were maintained in Dulbecco’s Modified Eagle Medium (DMEM) containing 10% FBS and 1% penicillin–streptomycin. All cells were incubated at 37°C in a humidified atmosphere with 5% CO_₂_.

### Reagents and Compounds

Curcumin analogues HO-3867 (Selleckchem, Cat. #S7501; CAS No. 1172133-28-6) and AKT-100 (from AKTX) were obtained from commercial vendors or synthesized as previously described (Smith et al., 2025; Williams et al., 2025). Hemin (Cat. #16009-13-5), zinc protoporphyrin IX (ZnPP) (Cat. # 15442-64-5), Liproxstatin-1 (Cat. #950455-15-9), and Z-VAD-FMK (Cat. # 187389-52-2) were all purchased from Sigma-Aldrich. Stock solutions were prepared in dimethyl sulfoxide (DMSO) and diluted in culture medium immediately before use to achieve the final treatment concentrations: HO-3867 (3–5 µM), AKT-100 (0.5–2 µM), Hemin (20–30 µM), ZnPP (5 µM), Liproxstatin-1 (50 nM), and Z-VAD-FMK (10 µM). A 1% DMSO solution was used as the vehicle control in all experiments.

### RNA Sequencing and Analyses

KLE, Hec50cc and Ishikawa cells were seeded in separate 6-well culture plates at 20,000 cells per well and allowed to adhere for 24 hours. At 24 hours post-seeding, cells were treated with either 0.1% DMSO (vehicle control) or AKT-100. After 24 hours of treatment, total RNA was extracted using the Qiagen RNeasy Mini Kit (Qiagen, Cat. #74104). RNA concentrations were measured using a Qubit Fluorometer. Ribosomal RNA was depleted using the RiboGone Kit, and cDNA libraries were constructed with the SMARTer Fragmentation Kit (Takara Bio, Cat. #634847 & Cat. #34938). Libraries were prepared using the Ion Plus Fragmentation Kit (Thermo Fisher, Cat. #4471252).

Sequencing was performed on Ion 550 chips using the S5XL Sequencing System. Sequencing reads were aligned to the human genome (RefSeq, UCSC hg19) using the RNA-Seq Alignment App (version 2.0.2) on BaseSpace (Illumina). Three biological replicates were included for differential gene expression analyses, comparing each treatment to the 0.1% DMSO control. Total RNA processing was conducted at the Analytical and Translational Genomics Core at the University of New Mexico. Genes with expression levels greater than 1 fragment per kilobase per million reads (FPKM) and showing greater than 2-fold changes were analyzed using Pathview Web (UNC Charlotte; https://pathview.uncc.edu/).

### Quantitative Reverse Transcription Polymerase Chain Reaction (RT-qPCR)

Cells were seeded in 6-well plates at a density of 20,000 cells per well and allowed to adhere overnight. KLE cells were treated with 0.1% DMSO, 2 µM AKT-100, 5 µM HO-3867, or 30 µM Hemin, whereas Hec50co cells were treated with 0.1% DMSO, 0.5 µM AKT-100, 3 µM HO-3867, or 20 µM Hemin. After 24 hours of treatment, total RNA was isolated using the RNeasy Mini Kit (Qiagen, Cat. #74104) following the manufacturer’s protocol. The curcumin analogue concentrations for these experiments were previously reported to be the IC50 values for these cell lines (Smith et al., 2025 and Williams et al., 2025)

Complementary DNA (cDNA) was synthesized from total RNA using the iScript cDNA Synthesis Kit (Bio-Rad). Quantitative PCR was performed using a SYBR Green detection system on a Applied Biosystems QuantStudio 3 Real-Time PCR System, 96-well (Thermo Fisher Scientific) with gene-specific primers. HMOX1 mRNA expression was normalized to 18S rRNA as an internal control, and relative expression levels were calculated using the 2^–ΔΔCt method. All experiments were performed in triplicate.

### Western Blotting

Cells were plated in 6-well plates at a density of 20,000 cells per well and allowed to attach overnight. KLE cells were treated with 0.1% DMSO, 2 µM AKT-100, 5 µM HO-3867, or 30 µM Hemin, whereas Hec50co cells were treated with 0.1% DMSO, 0.5 µM AKT-100, 3 µM HO-3867, or 20 µM Hemin. Cells were harvested and lysed in Pierce RIPA buffer supplemented with protease inhibitors, and protein concentrations were determined using the BCA Protein Assay Kit (Thermo Fisher Scientific). Equal amounts of protein (50 µg) were separated by SDS–PAGE on 10% polyacrylamide gels and transferred to nitrocellulose membranes. Membranes were blocked in 5% bovine serum albumin (BSA) and incubated overnight at 4°C with primary antibodies: HMOX1 (Proteintech, #66743), β-Actin (Santa Cruz, #sc-47778), or Tubulin (Proteintech, #66031), all at a 1:1000 dilution. After washing, membranes were incubated with HRP-conjugated anti-mouse or anti-rabbit secondary antibodies (1:1000 in 5% milk) for 1 hour at room temperature, followed by three washes in 1× TBST. Protein bands were visualized using enhanced chemiluminescence and quantified using ImageJ software. All experiments were performed in triplicate.

### Intracellular Iron and ROS Detection

Intracellular labile iron levels were assessed using FerroOrange (Dojindo Molecular Technologies,Inc., F374) following the manufacturer’s instructions. Cells were incubated with 1 µM FerroOrange for 45 minutes at 37 °C and visualized by fluorescence microscopy. Reactive oxygen species (ROS) levels were measured using 10 µM 2′,7′-dichlorofluorescin diacetate (DCFH-DA, Cayman, #85155). Cells were incubated with the probe for 45 minutes at 37 °C, washed with PBS, and fluorescence intensity was measured microscopically. All experiments were performed in triplicate, and fluorescence intensity was quantified using ImageJ software.

### Cell Viability and Proliferation Assay

For cell viability assays, KLE and Hec50co cells were seeded at an optimal density of 5,000 cells per well in 96-well plates with 100 µL of complete growth medium and allowed to adhere for 24 hours. After 24 hours, cells were treated with increasing concentrations of ZnPP, Liproxstatin-1, or Z-VAD-FMK in the presence of a constant concentration of AKT-100. Plates were incubated for 72 hours, and DNA content per well was quantified using the CyQUANT NF Cell Proliferation Assay (Thermo Fisher Scientific, Cat. #C35006). Fluorescence was measured on a Wallac 1420 Multilabel Counter (PerkinElmer). Treatments were performed in triplicate, and each experiment was independently repeated to confirm reproducibility.

### Gene Knockdown

KLE cells were seeded in 12-well plates (20,000 cells per well) and 96-well plates (5,000 cells per well) and allowed to attach overnight. HMOX1 Human Pre-designed siRNA Set A (MedChemExpress, #HY-RS06249) was combined with DreamFect Gold Transfection Reagent (OzBiosciences, #DG81000) and incubated for 5 minutes at room temperature. The mixture was then diluted in Opti-MEM I Reduced Serum Medium (Thermo Fisher Scientific, #51985034) and incubated for an additional 30 minutes to allow complex formation. The siRNA–transfection reagent mixture was added to complete growth medium to achieve a final siRNA concentration of 100 nM. For transfection, 1 mL of the siRNA–medium mixture was added to each well of a 12-well plate, and 100 µL was added to each well of a 96-well plate. Cells were incubated for 48 hours before treatment with 0.1% DMSO or 2 µM AKT-100.

After 24 hours of treatment, cells from the 12-well plates were collected; a portion was used for the colony formation assay, and the remainder was lysed for Western blot analysis to confirm gene knockdown efficiency. After 72 hours of treatment, cells in the 96-well plates were analyzed for cell viability using the CyQUANT NF Cell Proliferation Assay as described above.

### Colony Formation Assay

KLE cells (3,000 cells per well) obtained from the gene knockdown experiment were seeded in 6-well plates and cultured for two weeks to allow colony formation. Cells were washed once with phosphate-buffered saline (PBS) and fixed with 10% formaldehyde for 10 minutes at room temperature. After fixation, cells were stained with 0.05% crystal violet solution for 30 minutes, washed twice with tap water, and air-dried for 5 minutes. Bound crystal violet was solubilized in methanol, and absorbance was measured at 540 nm using a microplate reader to quantify colony formation.

### Statistical Analysis

All data are presented as mean ± standard deviation (SD). Statistical comparisons were performed using independent Student’s t-tests, one-way ANOVA, or two-way ANOVA, as appropriate. A p-value < 0.05 was considered statistically significant. All analyses were conducted using GraphPad Prism software.

## Results

### 1. Curcumin Analogues Modulate Ferroptosis-Related Gene Expression

To evaluate the effects of curcumin analogues on ferroptosis in endometrial cancer, three endometrial cancer cell lines (KLE, Hec50co and Ishikawa) were treated with HO-3867 or AKT-100 for 24 hours, followed by transcriptomic profiling using RNA sequencing. Pathway visualization using Pathview demonstrated activation of ferroptosis-related signaling, revealing coordinated increases in genes associated with iron uptake, intracellular iron handling, and oxidative stress responses. Conversely, ferroptosis-suppressive genes such as GPX4 were significantly downregulated following treatment with the curcumin analogues, reinforcing the pro-ferroptotic effects of these compounds (**Figure 1A–B**).

**Figure 1.**
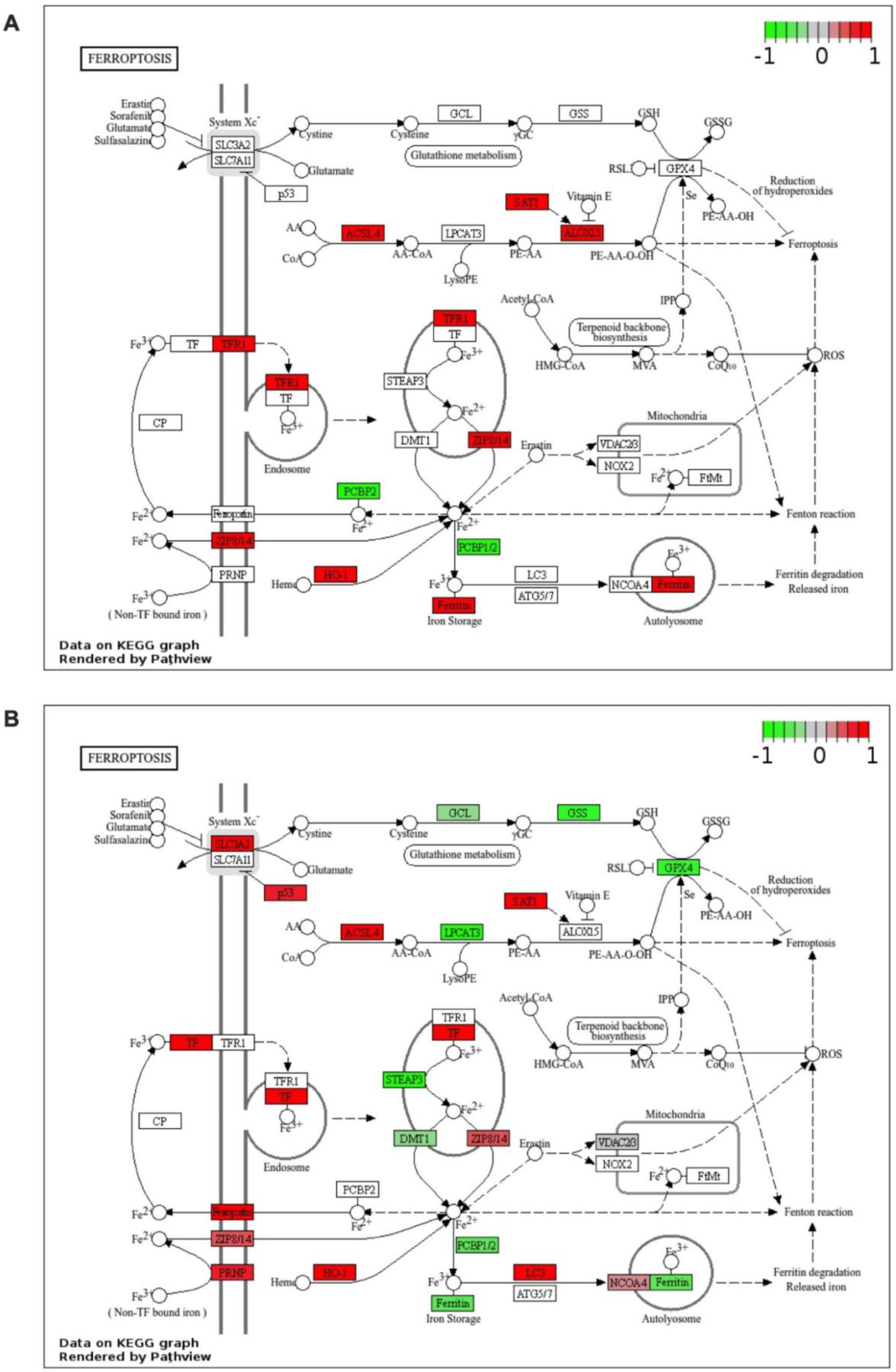
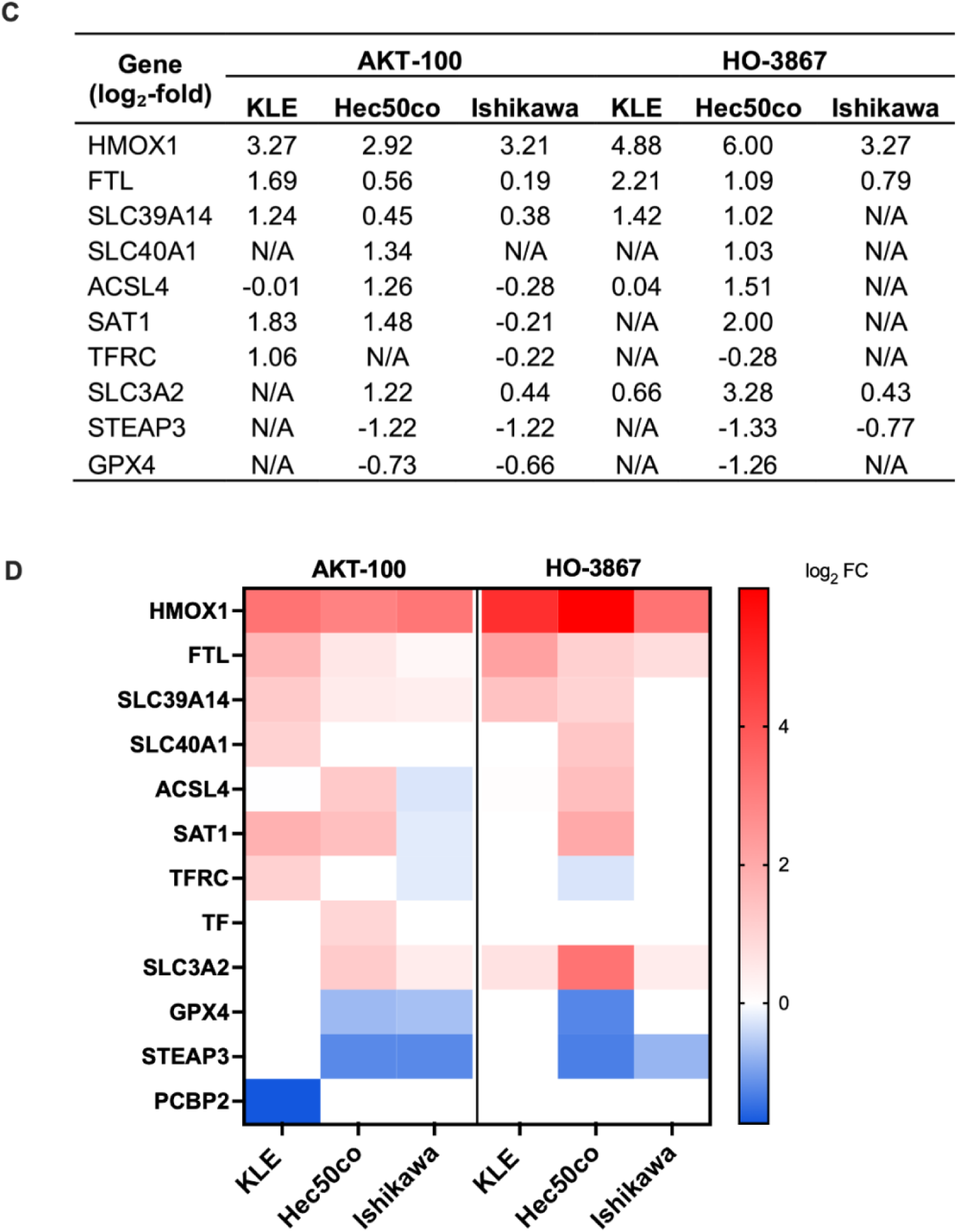
Ferroptosis-related gene expression changes revealed by RNA sequencing in endometrial cancer cells. Pathview visualization of differentially expressed genes in ferroptosis pathway in (A) KLE cells treated with 2 µM AKT-100 for 24 hours and (B) Hec50co cells treated with 0.5 µM AKT-100 for 24 hours. Red indicates upregulation and green indicates downregulation relative to the 0.1% DMSO control. (C) Table summarizing the differentially expressed genes (log₂-fold change) among three endometrial cell lines after 24h treatment with AKT-100, and HO-3867 compared to 0.1% DMSO control. (D) Heatmap further displaying the upregulated (red) and downregulated (blue) genes.

Heme oxygenase-1 (HMOX1) exhibited the most pronounced upregulation across all three cell lines following 24-hour treatment with either analogue. In KLE cells, 2 µM AKT-100 increased HMOX1 expression by 4.88 log₂-fold, whereas 5 µM HO-3867 induced a 3.27 log₂-fold increase. In Hec50co cells, treatment with 0.5 µM AKT-100 and 3 µM HO-3867 produced 2.92 and 5.99 log₂-fold increases, respectively. In Ishikawa cells, 0.5 µM AKT-100 and 2 µM HO-3867 led to 3.21 and 3.27 log₂-fold increases, respectively (**Figure 1C–D**). Beyond HMOX1, several genes involved in iron transport and storage—including SLC39A14 (ZIP14), SLC39A8 (ZIP8), and FTL (ferritin light chain)—were consistently upregulated across all three cell lines (**Figure 1C–D**).

To expand these findings to ovarian cancer, additional RNA-seq analyses from ovarian cancer cell lines (OVCAR3 and COV362) were performed. These datasets similarly demonstrate that both AKT-100 and HO-3867 induce similar robust ferroptosis-related transcriptional changes (**Figure S1**)—highlighting the broader applicability of curcumin analogues in promoting ferroptotic cell death across multiple gynecologic cancer models.

### 2. Upregulation of HMOX1 Confirmed by qPCR and Immunoblotting

To validate the RNA-seq findings, HMOX1 mRNA and protein levels were assessed using real-time quantitative PCR (RT-QPCR) and immunoblotting. Both AKT-100 and HO-3867 significantly increased HMOX1 gene expression in KLE (**Figure 2A**) and Hec50co cells (**Figure 2B**) following 24-hour treatments, exhibiting patterns comparable to the known HMOX1 inducer, hemin. Immunoblot analyses further confirmed a marked increase in HMOX1 protein levels in both cell lines (**Figure 2C–D**), supporting the conclusion that curcumin analogues transcriptionally and translationally activate HMOX1 expression.

**Figure 2.**
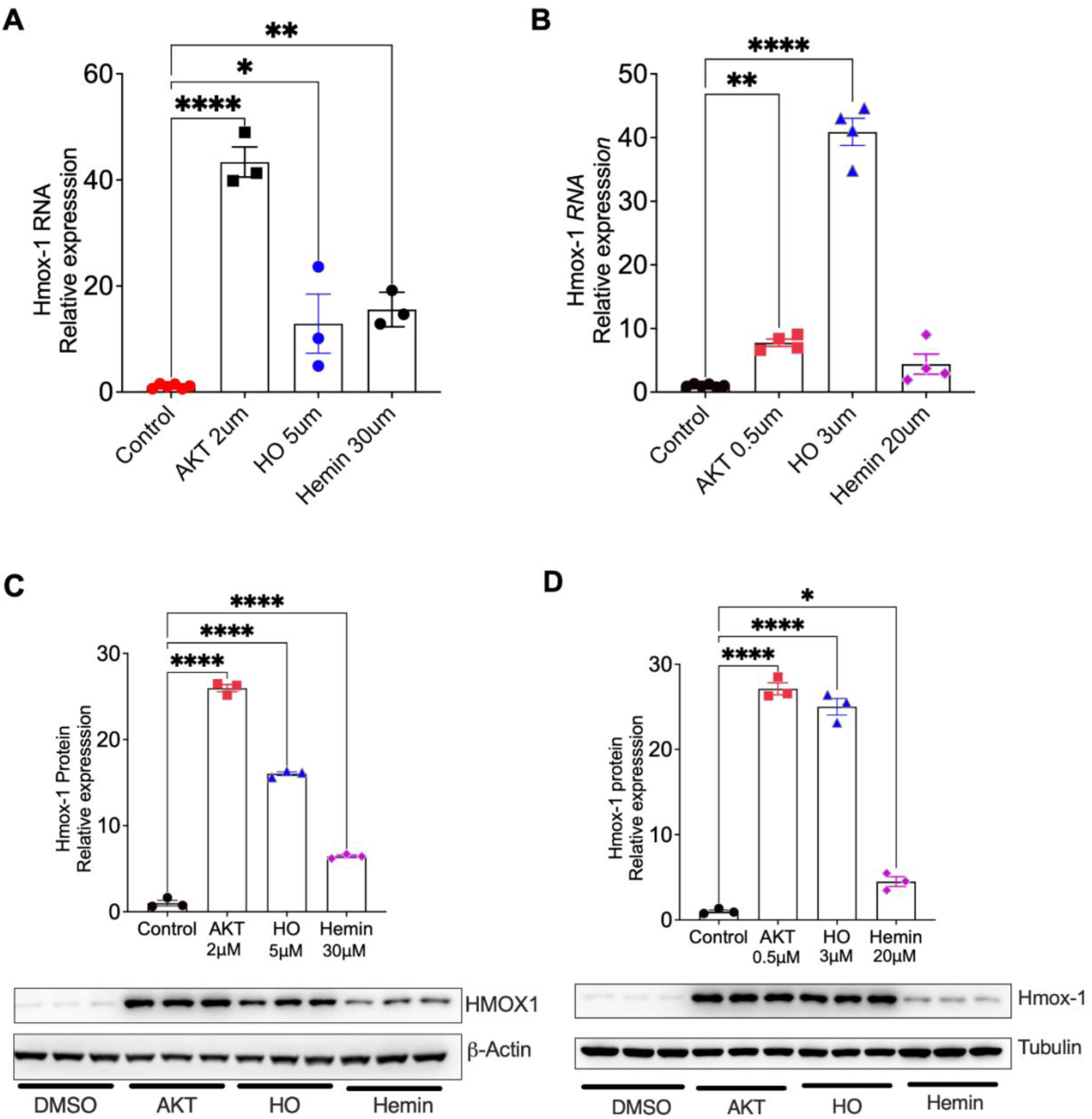
Curcumin analogues significantly elevate *HMOX1* transcription and translation in endometrial cancer cells. (A) Real-time PCR analysis of *HMOX1* mRNA expression in KLE cells treated with 2 µM AKT-100, 5 µM HO-3867, or 30 µM hemin for 24 hours. (B) Real-time PCR analysis of *HMOX1* mRNA expression in Hec50co cells treated with 0.5 µM AKT-100, 3 µM HO-3867, or 20 µM hemin for 24 hours. (C) Immunoblotting and densitometric quantification of HMOX1 protein levels in KLE cells under the same treatment conditions. (D) Immunoblotting and densitometric quantification of HMOX1 protein levels in Hec50co cells under the same treatment conditions. Data are presented as mean ± SD. *p* < 0.05, *p* < 0.01, *p* < 0.001; one-way ANOVA followed by Dunnett’s multiple comparisons test.

### 3. AKT-100 increases intracellular iron and ROS levels

Given the established role of HMOX1 in regulating iron mobilization and oxidative stress (**Figure 3A**), we next evaluated whether AKT-100 treatment alters intracellular iron and ROS levels in KLE cells. Treatment with 2 µM AKT-100 for 24 hours markedly increased labile iron, as visualized by FerroOrange staining (**Figure 3B**). In parallel, total ROS levels, detected using the fluorescent probe DCFH-DA, were significantly elevated compared with vehicle-treated controls (**Figure 3C**). These results indicate that AKT-100 enhances intracellular iron accumulation and oxidative stress, consistent with ferroptotic activation.

**Figure 3.**
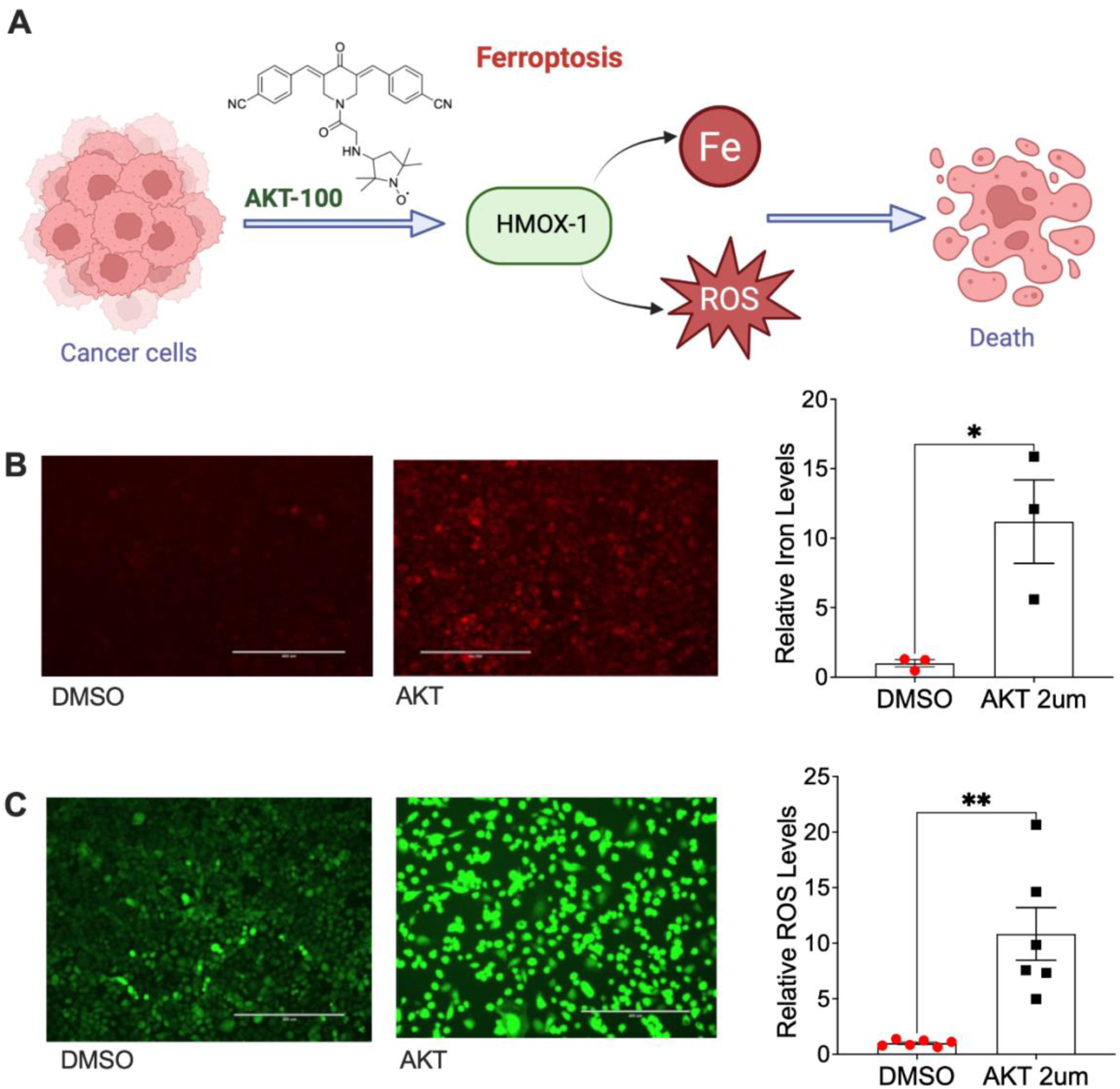
AKT-100 significantly increases intracellular iron and ROS levels in KLE cells. (A) Graphical illustration showing how novel curcumin analogues induce ferroptosis via heme oxygenase-1 (HMOX1) upregulation to suppress endometrial cancer growth. (B) FerroOrange staining assessing intracellular iron levels in KLE cells treated with 2 µM AKT-100 for 24 hours. (C) DCFH-DA staining measuring total ROS levels under the same treatment conditions. Data are presented as mean ± SD. *p* < 0.05, *p* < 0.01; unpaired t-test.

### 4. AKT-100 Induces Cell Death Through HMOX1–Mediated Ferroptosis and Apoptosis

To directly assess whether HMOX1–mediated ferroptosis contributes to the cytotoxic effects of AKT-100, KLE cells were co-treated with the HMOX1 inhibitor zinc protoporphyrin IX (ZnPP) or the ferroptosis inhibitor liproxstatin-1. CyQUANT proliferation assays demonstrated that both ZnPP (5 µM) and liproxstatin-1 (50 nM) partially reversed the growth-inhibitory effects induced by 2 µM AKT-100 (**Figure 4A–B**). To further elucidate the mechanisms underlying AKT-100–induced cell death, KLE cells were additionally co-treated with the pan-caspase inhibitor Z-VAD-FMK (10 µM). Treatment with Z-VAD-FMK partially attenuated the cytotoxic effects of AKT-100 (**Figure 4C**), indicating that both apoptosis and ferroptosis contribute to AKT-100–mediated cell death, consistent with previous reports (Williams et al., 2025).

**Figure 4.**
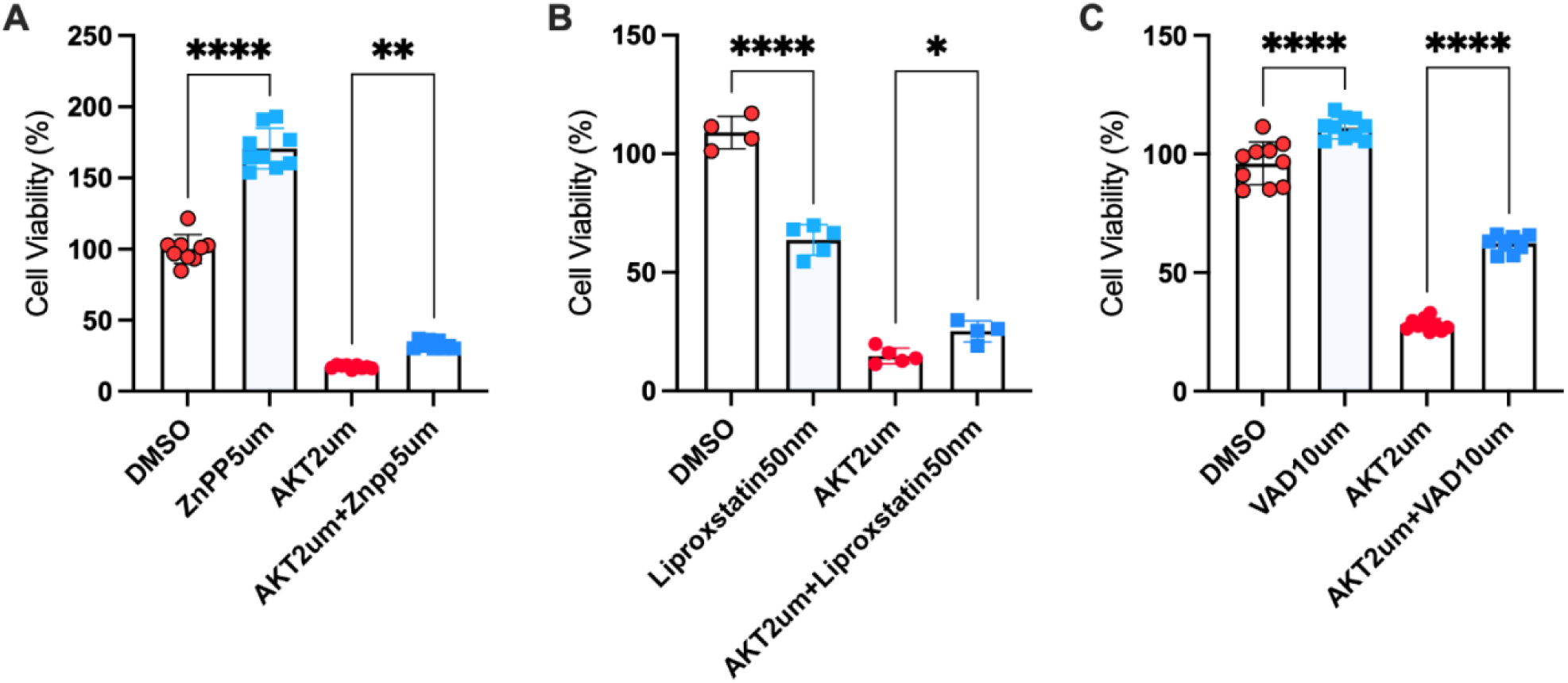
HMOX1, ferroptosis, and apoptosis inhibition partially reverse AKT-100–induced cytotoxicity in KLE cells. CyQUANT cell proliferation assays of KLE cells treated with 2 µM AKT-100 in combination with: (A) 5 µM zinc protoporphyrin IX (ZnPP, HMOX1 inhibitor), (B) 50 nM liproxstatin-1 (ferroptosis inhibitor), or (C) 10 µM Z-VAD-FMK (pan-caspase/apoptosis inhibitor) for 72 hours. Data are presented as mean ± SD. *p* < 0.05, \**p* < 0.01, \*\*\**p* < 0.0001; two-way ANOVA followed by Šídák’s multiple comparisons test.

Collectively, these findings indicate that HMOX1 induction and subsequent ferroptosis signaling contribute, at least in part, to the anti-proliferative activity of AKT-100 in endometrial cancer cells. The concurrent activation of ferroptosis and apoptosis further suggests that AKT-100 may overcome resistance mechanisms typically associated with apoptosis-targeted therapies.

### 5. Suppression of HMOX1 Expression via siRNA Attenuates AKT-100–Induced Toxicity and Promotes Cell Growth

To investigate the role of HMOX1 in mediating AKT-100–induced cytotoxicity, siRNA-mediated knockdown experiments were performed in KLE cells. Immunoblot analysis confirmed that transfection with HMOX1–specific siRNA substantially reduced HMOX1 protein levels compared with negative control siRNA (**Figure 5A**). Silencing HMOX1 significantly mitigated the reduction in cell viability caused by AKT-100 treatment, as assessed by CyQUANT proliferation assays (**Figure 5B**). Consistently, HMOX1 knockdown enhanced the clonogenic capacity of AKT-100–treated KLE cells (**Figure 5C**), and quantitative analysis of colony formation corroborated this effect (**Figure 5D**). Together, these findings indicate that suppression of HMOX1 expression attenuates AKT-100–induced cytotoxicity and promotes cell growth, suggesting that HMOX1 plays a critical role in the antiproliferative activity of AKT-100 in endometrial cancer cells.

**Figure 5.**
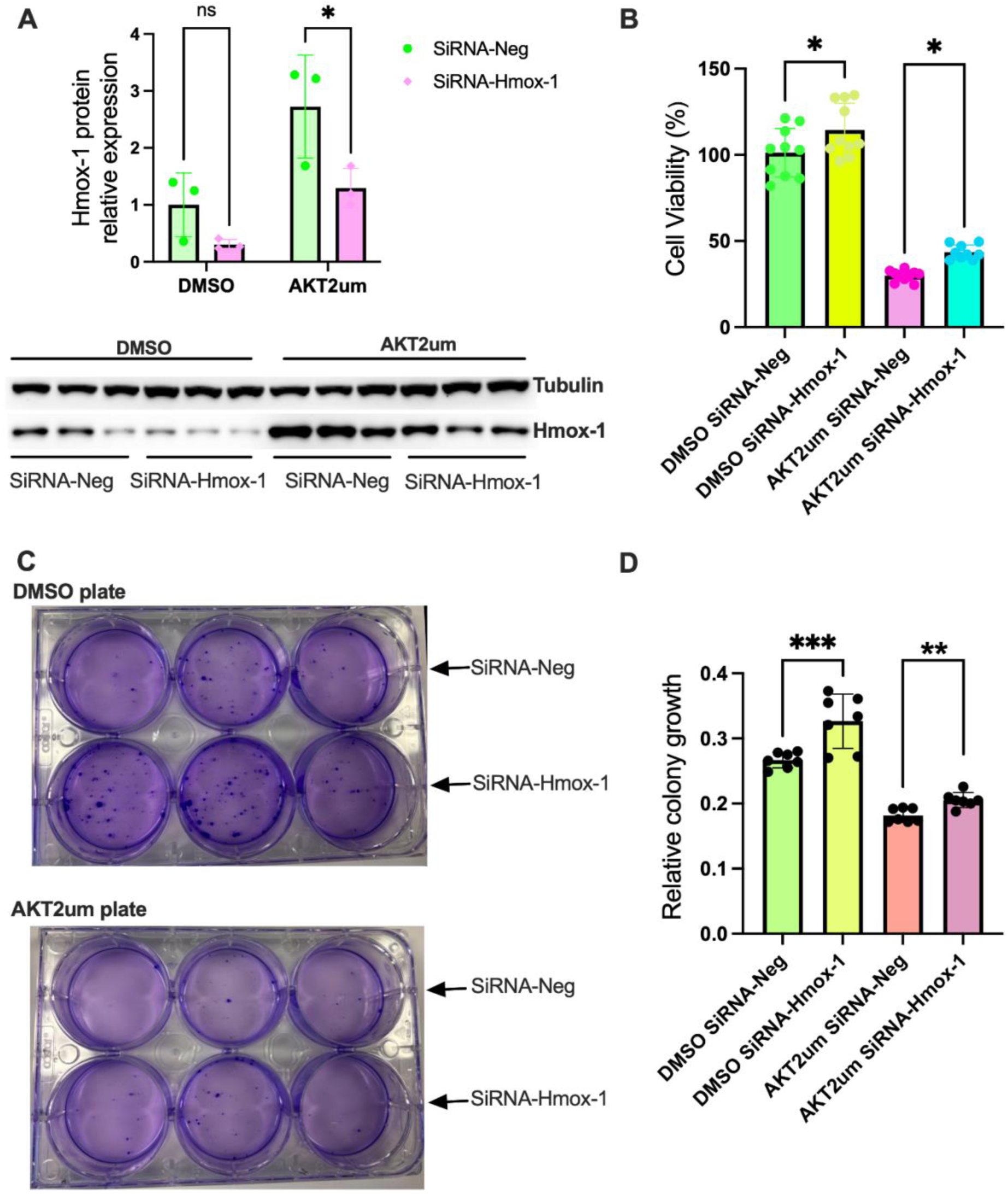
Knockdown of HMOX1 via siRNA attenuates AKT-100–induced cytotoxicity and promotes cell growth in KLE cells. (A) Immunoblotting and densitometric quantification of HMOX1 protein levels in KLE cells transfected with 100 nM negative control siRNA (siRNA-Neg) or HMOX-1–specific siRNA (siRNA-HMOX1) for 48 hours, followed by treatment with 2 µM AKT-100 for 24 hours. Data were analyzed using two-way ANOVA followed by Šídák’s multiple comparisons test. (B) CyQUANT cell proliferation assay of KLE cells transfected with 100 nM siRNA for 48 hours and then treated with 2 µM AKT-100 for 72 hours. Data were analyzed using one-way ANOVA followed by Tukey’s multiple comparisons test. (C) Representative images of colony formation in KLE cells transfected with 100 nM siRNA for 48 hours and treated with 2 µM AKT-100 for 24 hours. (D) Quantification of colony formation shown in (C). Data were analyzed using one-way ANOVA followed by Tukey’s multiple comparisons test. Data are presented as mean ± SD. *p* < 0.05, \**p* < 0.01, \*\**p* < 0.001; ns, not significant.

## Discussion

In this study, we demonstrate that the bioavailable curcumin analogues HO-3867 and AKT-100 robustly induce ferroptosis-associated gene expression in endometrial cancer cells, with a pronounced upregulation of heme oxygenase-1 (HMOX1). RNA sequencing, RT-qPCR, and immunoblotting consistently revealed that treatment with these compounds strongly elevates HMOX1, along with genes involved in iron transport and storage, including SLC39A14 (ZIP14), SLC39A8 (ZIP8), and ferritin light chain (FTL). Functionally, this transcriptional program is accompanied by increased intracellular iron and ROS levels—hallmarks of ferroptosis. Notably, pharmacological inhibition of HMOX1 with zinc protoporphyrin IX (ZnPP) or blockade of ferroptosis with liproxstatin-1 partially rescued AKT-100–induced cytotoxicity, highlighting the central role of HMOX1–mediated ferroptosis in the anticancer activity of these analogues. Furthermore, inhibition of apoptosis with Z-VAD-FMK also attenuated AKT-100–induced cell death, indicating that this curcumin analogue can trigger apoptosis in addition to ferroptosis.

Although curcumin itself has been reported to induce ferroptosis in various tumor types, its clinical application is limited by poor bioavailability (Xin & Zhang, 2024). Our findings demonstrate that optimized curcumin analogues such as HO-3867 and AKT-100 retain curcumin’s ferroptosis-inducing potential while exhibiting activity at low micromolar to nanomolar concentrations (Smith et al., 2025; Williams et al., 2025). This is particularly relevant in endometrial cancer, where conventional chemotherapies often display suboptimal efficacy (Brasseur et al., 2022). By concurrently activating both ferroptotic and apoptotic pathways, these compounds provide a dual mechanism of tumor suppression that may overcome resistance to apoptosis-targeted therapies.

Our results also reinforce the emerging role of HMOX1 as a critical regulator of ferroptosis. HMOX1 catalyzes heme degradation into biliverdin, carbon monoxide, and free iron, thereby increasing the labile iron pool and promoting ROS accumulation (Ryter & Tyrrell, 2021). Here, induction of HMOX1 by curcumin analogues establishes a ferroptotic environment selectively toxic to endometrial cancer cells. The reversal of cytotoxicity by ZnPP further supports the mechanistic contribution of HMOX1 in this process. These findings suggest that HMOX1 expression may serve as a predictive biomarker for response to ferroptosis-inducing therapies in endometrial cancer.

Ferroptosis-based therapies are gaining attention as promising strategies to overcome drug resistance, particularly in cancers that evade apoptosis-driven treatments (Zhang et al., 2022). Our data provide preclinical evidence supporting HO-3867 and AKT-100 as strong candidates for the treatment of advanced or recurrent endometrial cancer. Notably, our supplemental RNA-seq data from ovarian cancer cell lines suggest that the pro-ferroptotic effects of these analogues are not limited to endometrial cancer, supporting broader applicability in gynecologic malignancies.

The reported low toxicity of HO-3867 in non-cancerous cells (Selvendiran et al., 2010) further underscores their translational potential. Ferroptosis inducers may synergize with existing therapies, including chemotherapy, radiotherapy, or immune checkpoint inhibitors, all of which modulate tumor oxidative stress and iron metabolism (Jiang et al., 2021). Our team found that HO-3867 is synergistic with the PARP inhibitor olaparib in both endometrial and ovarian cancer cell lines and further limited the tumor growth in the OVCAR3 xenograft mouse model. (Smith et al., 2025)

Despite these promising results, certain limitations warrant consideration. Our current findings are predominantly based on *in vitro* and xenograft models; comprehensive pharmacokinetic, safety, and efficacy studies in more physiologically relevant *in vivo* models are necessary to validate clinical potential. Additionally, while HMOX1 appears to be a key mediator of ferroptosis induction, the contribution of other ferroptosis-regulating pathways, including GPX4 (Xue et al., 2024), lipid peroxidation enzymes, and iron-handling proteins, remains to be fully elucidated. Future studies dissecting these interactions will provide a more complete understanding of curcumin analogue–induced ferroptosis and may identify additional biomarkers or combinatorial targets.

In conclusion, our study demonstrates that HO-3867 and AKT-100 are potent inducers of HMOX1-mediated ferroptosis in endometrial cancer, with concurrent activation of apoptotic pathways. These bioavailable curcumin analogues effectively disrupt iron homeostasis and oxidative stress regulation, leading to selective tumor cell death. By integrating transcriptional, biochemical, and functional evidence, our work provides a compelling preclinical rationale for targeting HMOX1–regulated ferroptosis as a therapeutic strategy in advanced and drug-resistant endometrial cancer, and potentially other gynecologic malignancies. These findings support further development of HO-3867 and AKT-100 as novel anticancer agents, either as monotherapies or in combination with existing treatments, to improve outcomes in patients with refractory disease.

## Acknowledgments

X.X. was partly supported by the National Institutes of Health (1R01ES035780, P50CA265793), John f. Perkins, Jr. Research Career Enhancement Award from the American Physiological Society, the HSC Pilot Funding Program from the University of New Mexico (UNM) HSC Office of Research, a Research Pilot Project Grant from University of New Mexico Environmental Health Signature Program and Superfund (P42 ES025589), an Off-Setting Cuts Pilot Project from UNM comprehensive cancer center (UNMCCC, P30CA118100), Health Sciences & Main Campus Research Collaboration Seed Grant Award from UNM Rainforest Innovations. X. W. was partially supported by a Matching Support Pilot Project Award from UNM comprehensive cancer center (No. P30CA118100). K.K.L. was supported by the following grants: NIH 5R01CA99908–22, CDMRP CA210610, CDMRP CA2220729, NIH P50CA264793, NIH P01CA872735, and NIH P30CA118100, with support from the UNMCCC Biostatistics and the ATG Genomics Shared Resources.

## References

Arcos, M., Goodla, L., Kim, H., Desai, S. P., Liu, R., Yin, K., Liu, Z., Martin, D.R., and Xue, X. (2024). PINK1-deficiency facilitates mitochondrial iron accumulation and colon tumorigenesis. Autophagy. 10.1080/15548627.2024.2425594.

Brasseur K, Gévry N, Asselin E. Chemoresistance and targeted therapies in ovarian and endometrialcancers. Oncotarget. 2017 Jan 17;8(3):4008–4042. doi: 10.18632/oncotarget.14021.

Bristow, R.E., Zerbe, M.J., Rosenshein, N.B., Grumbine, F.C., and Montz, F.J. (2000). Stage IVB Endometrial Carcinoma: The Role of Cytoreductive Surgery and Determinants of Survival. Gynecol. Oncol. 78, 85–91.

Carlson M, Thiel K, Leslie K. Past, present, and future of hormonal therapy in recurrent endometrial cancer. Int J Womens Health. 2014; 6:429–435.

Datla, S.R., Dusting, G.J., Mori, T.A., Taylor, C.J., Croft, K.D., and Jiang, F. (2007). Induction of Heme Oxygenase-1 In Vivo Suppresses NADPH Oxidase–Derived Oxidative Stress. Hypertension 50, 636–642.

Dixon, S.J., Lemberg, K.M., Lamprecht, M.R., Skouta, R., Zaitsev, E.M., Gleason, C.E., Patel, D.N., Bauer, A.J., Cantley, A.M., Yang, W.S., et al. (2012). Ferroptosis: An Iron-Dependent Form of Nonapoptotic Cell Death. Cell 149, 1060–1072.

Friedmann Angeli, J.P., Krysko, D. V, and Conrad, M. (2019). Ferroptosis at the crossroads of cancer-acquired drug resistance and immune evasion. Nat. Rev. Cancer 19, 405–414.

Ghoochani A, Hsu EC, Aslan M, Rice MA, Nguyen HM, Brooks JD, Corey E, Paulmurugan R, Stoyanova T. Ferroptosis Inducers Are a Novel Therapeutic Approach for Advanced Prostate Cancer. Cancer Res. 2021 Mar 15;81(6):1583–1594. doi: 10.1158/0008-5472.CAN-20-3477. Epub 2021 Jan 22.PMID: 33483372

Hsueh K-C, Ju P-C, Hsieh Y-H, Su S-C, Yeh C-B, Lin C-W. HO-3867, a curcumin analog, elicits cell apoptosis and p38-mediated caspase activation in hepatocellular carcinoma. Environmental Toxicology. 2024; 39(2): 794–802.

Kim, H., Villareal, L.B., Liu, Z., Haneef, M., Falcon, D.M., Martin, D.R., Lee, H.-J., Dame, M.K., Attili, D., Chen, Y., et al. (2023). Transferrin Receptor-Mediated Iron Uptake Promotes Colon Tumorigenesis. Adv. Sci. n/a, 2207693. 10.1002/advs.202207693.

Li Z, Caron de Fromentel C, Kim W, Wang WH, Sun J, Yan B, Utturkar S, Lanman NA, Elzey BD, Yeo Y, Zhang H, Kazemian M, Levrero M, Andrisani O. RNA helicase DDX5 modulates sorafenib sensitivity in hepatocellular carcinoma via the Wnt/beta-catenin-ferroptosis axis. Cell Death Dis. 2023 Nov 30;14(11):786. doi: 10.1038/s41419-023-06302-0.

Lin, H., Chen, X., Zhang, C., Yang, T., Deng, Z., Song, Y., Huang, L., Li, F., Li, Q., Lin, S., et al. (2021). EF24 induces ferroptosis in osteosarcoma cells through HMOX1. Biomed. Pharmacother. 136, 111202.

Liu, S., Zhang, Q., Liu, W., and Huang, X. (2022). Prediction of Prognosis in Patients With Endometrial Carcinoma and Immune Microenvironment Estimation Based on Ferroptosis-Related Genes. DOI=10.3389/fmolb.2022.916689.

Lu, P.W.-A.; Chou, C.-H.; Yang, J.-S.; Hsieh, Y.-H.; Tsai, M.-Y.; Lu, K.-H.; Yang, S.-F. HO-3867 Induces Apoptosis via the JNK Signaling Pathway in Human Osteosarcoma Cells. Pharmaceutics 2022, 14, 1257.

Mao C, Jiang D, Koong AC, Gan B. Exploiting metabolic cell death for cancer therapy. Nat Rev Cancer. 2025 Oct 15. doi: 10.1038/s41568-025-00879-8. Online ahead of print.PMID: 41094043

McEachron, J., Chatterton, C., Hastings, V., Gorelick, C., Economos, K., Lee, Y.-C., and Kanis, M.J. (2020). A clinicopathologic study of endometrial cancer metastatic to bone: Identification of microsatellite instability improves treatment strategies. Gynecol. Oncol. Reports 32, 100549. 10.1016/j.gore.2020.100549.

Mohammad RM, Muqbil I, Lowe L, Yedjou C, Hsu HY, Lin LT, Siegelin MD, Fimognari C, Kumar NB, Dou QP, Yang H et al. Broad targeting of resistance to apoptosis in cancer. Semin Cancer Biol. 2015 Dec;35 Suppl(0):S78–S103. doi:10.1016/j.semcancer.2015.03.001. Epub 2015 Apr 28. PMID: 25936818

Rath KS, Naidu SK, Lata P, Bid HK, Rivera BK, McCann GA, Tierney BJ, Elnaggar AC, Bravo V, Leone G, et al. HO-3867, a safe STAT3 inhibitor, is selectively cytotoxic to ovarian cancer. Cancer Res. 2014 Apr 15;74(8):2316–27. doi: 10.1158/0008-5472.CAN-13-2433. Epub 2014 Mar 3. PMID: 24590057

Schwartz, A.J., Goyert, J.W., Solanki, S., Kerk, S.A., Chen, B., Castillo, C., Hsu, P.P., Do, B.T., Singhal, R., Dame, M.K., et al. (2021). Hepcidin sequesters iron to sustain nucleotide metabolism and mitochondrial function in colorectal cancer epithelial cells. Nat. Metab. 3, 969–982. 10.1038/s42255-021-00406-7.

Selvendiran, K., Ahmed, S., Dayton, A., Kuppusamy, M.L., Tazi, M., Bratasz, A., Tong, L., Rivera, B.K., Kálai, T., Hideg, K., et al. (2010). Safe and targeted anticancer efficacy of a novel class of antioxidant-conjugated difluorodiarylidenyl piperidones: Differential cytotoxicity in healthy and cancer cells. Free Radic. Biol. Med. 48, 1228–1235.

Siegel, R.L., Miller, K.D., Fuchs, H.E., and Jemal, A. (2022). Cancer statistics, 2022. CA. Cancer J. Clin.72, 7–33.

Smith LE, Padilla JL, Licor A, Steinkamp MP, Lagutina IV, Guo Y, Devor EJ, Pankratz VS, Sallmyr A, Oyebamiji OM, Choe JY, Williams GL, Leslie KK. Novel p53 reactivators that are synergistic with olaparib for the treatment of gynecologic cancers with mutant p53. Transl Oncol. 2025 Nov;61:102522. doi:10.1016/j.tranon.2025.102522. Epub 2025 Sep 6.PMID: 40915175

Soares MP, Bach FH. Heme oxygenase-1: from biology to therapeutic potential. Trends Mol Med. 2009 Feb;15(2):50–8. doi: 10.1016/j.molmed.2008.12.004. Epub 2009 Jan 21.PMID: 19162549

Williams, G.L., Smith, L.E., Padilla, J.L., Goss, A.S., Ortega, D.B., Wu, X., Daw, J.N., Mitra, S., Jagtap, P., Choe, J., Leslie, K.K. The novel curcumin analogue AKT-100 targets mutant p53 in gynecologic cancer cells bioRxiv 2025.10.06.680794; doi: 10.1101/2025.10.06.680794

Wu, J., Zhang, L., Wu, S., and Liu, Z. (2022). Ferroptosis: Opportunities and Challenges in Treating Endometrial Cancer. Biosci. 9, 929832.

Wu, L., Xu, G., Li, N., Zhu, L., and Shao, G. (2023). Curcumin Analog, HO-3867, Induces Both Apoptosis and Ferroptosis via Multiple Mechanisms in NSCLC Cells with Wild-Type p53. Evidence-Based Complement. Altern. Med. 2023, 8378581.

Xie, Y., Hou, W., Song, X., Yu, Y., Huang, J., Sun, X., Kang, R., and Tang, D. (2016). Ferroptosis: process and function. Cell Death Differ. 23, 369–379.

Xin, W., and Zhang, Y. (2024). Curcumin activates the JNK signaling pathway to promote ferroptosis in colon cancer cells. Chem. Biol. Drug Des. 103, e14468. 10.1111/cbdd.14468.

Xue, X., Bredell, B.X., Anderson, E.R., Martin, A., Mays, C., Nagao-Kitamoto, H., Huang, S., Győrffy, B., Greenson, J.K., Hardiman, K., et al. (2017). Quantitative proteomics identifies STEAP4 as a critical regulator of mitochondrial dysfunction linking inflammation and colon cancer. Proc. Natl. Acad. Sci. 114, E9608–E9617. 10.1073/pnas.1712946114.

Xue X, Liu Z, Liang Y, Kwon YY, Liu R, Martin D, Hui S. Glutathione peroxidase 4 suppresses manganese-dependent oxidative stress to reduce colorectal tumorigenesis. Res Sq [Preprint]. 2024 Jan 8: rs.3.rs-3837925. doi: 10.21203/rs.3.rs-3837925/v1. PMID: 38260380; PMCID: PMC10802749.

Xue, X., Ramakrishnan, S.K., Weisz, K., Triner, D., Xie, L., Attili, D., Pant, A., Győrffy, B., Zhan, M., Carter-Su, C., et al. (2016). Iron Uptake via DMT1 Integrates Cell Cycle with JAK-STAT3 Signaling to Promote Colorectal Tumorigenesis. Cell Metab. 24, 447–461. 10.1016/j.cmet.2016.07.015.

Xue, X., Taylor, M., Anderson, E., Hao, C., Qu, A., Greenson, J.K., Zimmermann, E.M., Gonzalez, F.J., and Shah, Y.M. (2012). Hypoxia-Inducible Factor-2α Activation Promotes Colorectal Cancer Progression by Dysregulating Iron Homeostasis. Cancer Res. 72, 2285–22

Zhang C, Liu X, Jin S, Chen Y, Guo R. Ferroptosis in cancer therapy: a novel approach to reversing drug resistance. Mol Cancer. 2022 Feb 12;21(1):47. doi: 10.1186/s12943-022-01530-y.

Zuo L, Zou X, Ge J, Hu S, Fang Y, Xu Y, Chen R, Xu S, Yu G, Zhou X, Ji L. The Nrf2-HMOX1 pathway as a therapeutic target for reversing cisplatin resistance in non-small cell lung cancer via inhibiting ferroptosis. Cell Death Discov. 2025 Jun 21;11(1):287. doi: 10.1038/s41420-025-02564-z.PMID: 40544

